# Synthetic Hibernation Suppresses Osteosarcoma Growth and Protects Normal Tissue from Radiation

**DOI:** 10.64898/2026.11.11.764778

**Authors:** Reema Chowdhury, Emiliana Piscitiello, Tamara Vitacchio, Olga Sokol, Malte Benje, Julius Oppermann, Alexander Helm, Niccolo E. Saccone, Gaia Volpi, Nivida Shete, Jian Xian Teh, Anggraeini Puspitasari Kokko, Marco Durante, Matteo Cerri, Walter Tinganelli

**Author notes:** These authors contributed equally to this work and share first authorship (Reema Chowdhury, Emiliana Piscitiello). These authors contributed equally to this work and share senior authorship (Matteo Cerri, Walter Tinganelli).

## Abstract

Natural hibernation can arrest tumor growth, but it remains unclear whether, and to what extent, repeated pharmacologically induced synthetic torpor (ST) can reproduce this effect. It is also unknown whether the radioprotection conferred by natural and synthetic torpor extends to tumors, potentially compromising radiotherapy.

Here, five 5′-AMP–induced torpor bouts administered every other day, a protocol designed to mimic repeated hibernation cycles (synthetic hibernation), markedly restrained osteosarcoma growth in female C3H/HeNRj mice bearing syngeneic LM8 tumors. Interestingly, growth suppression persisted long after the final torpor bout. When 6 Gy total-body X-ray irradiation was delivered during the second bout, tumor growth control was comparable with irradiation under normothermia, whereas ovarian follicular atresia was lower and remained similar to non-irradiated controls. Together, these findings support synthetic hibernation as a strategy to restrain tumor growth and mitigate normal-tissue radiation injury without compromising tumor control.

## INTRODUCTION

Uncontrolled cell proliferation is a hallmark of cancer, a disease that remains a leading cause of death worldwide (*1, 2*). Natural hibernation offers a model for exploring how this proliferation might be restrained. This survival strategy is characterized by multiday bouts of torpor, a period of profound, reversible hypometabolism and hypothermia, which allow animals, including primates (*3, 4*), to endure adverse environmental conditions (*3–5*).

Central to this hypometabolic state is a coordinated suppression of cellular activity, including cell-cycle arrest (*6, 7*). Notably, this suppression seems to extend to malignant cells, which appear particularly sensitive to hypothermic conditions (*8–11*). Evidence for this phenomenon stems mainly from classical studies in hibernators, where tumor growth was found to halt during the hibernation phase and resume only upon return to euthermia (*12, 13*). Despite these early findings, this specific line of research was long left unpursued.

Today, artificially induced torpor or synthetic torpor (ST) is emerging as a powerful translational tool, and multiple procedures (*14-17*) have been proposed to mimic significant features of torpor (*18-21*). Among these, the increase in cell radioprotection has raised interest in both space and medical research (*22, 24*). Our group previously demonstrated that pharmacologically induced ST achieved either via central RPa inhibition or administering a single dose of 5′-AMP (*25, 26*) confers radioprotection in non-hibernating animals.

Following earlier works on induced hypothermia in tumor-bearing mice (27), recent studies suggest that induced torpor-like states may also slow tumor growth (*28-31*). However, the mechanisms and experimental conditions underlying this tumor-suppressive effect remain poorly characterized. Crucially, whether tumor tissue itself acquires radioresistance during ST, which would inherently compromise radiotherapy efficacy, has not yet been investigated.

In this study, we tested whether repeated bouts of pharmacologically induced ST, an approach we term “synthetic hibernation” (SH), suppress tumor progression in osteosarcoma-bearing mice. SH was used to better approximate the sustained tumor-suppressive state observed during natural hibernation. We then combined SH with radiotherapy to address two questions central to its translational potential: whether SH confers systemic radioprotection and, critically, whether this protection extends to the tumor tissue itself.

## RESULTS

### Repeated 5’-AMP administration produces a reproducible and well tolerated SH

All animals completed the five-bout protocol, and no procedure-related adverse effects were observed. Each 5’-AMP injection produced a reproducible reduction in rectal temperature, although bout duration varied among animals. During the 4-h monitoring window, temperature fell by 9.92 ± 0.74 °C in experiment 1 and 9.38 ± 0.59 °C in experiment 2. Mean nadir temperatures were 24.63 ± 0.80 °C and 24.17 ± 0.59 °C in experiments 1 and 2, respectively. At 4 h post-injection, rectal temperature remained below baseline (32.45 ± 0.47 °C in experiment 1; 30.87 ± 0.30 °C in experiment 2) (Fig. 1B, 2B), indicating partial rather than complete rewarming within the monitoring window. Data are expressed as mean ± SEM.

**Figure 1.**
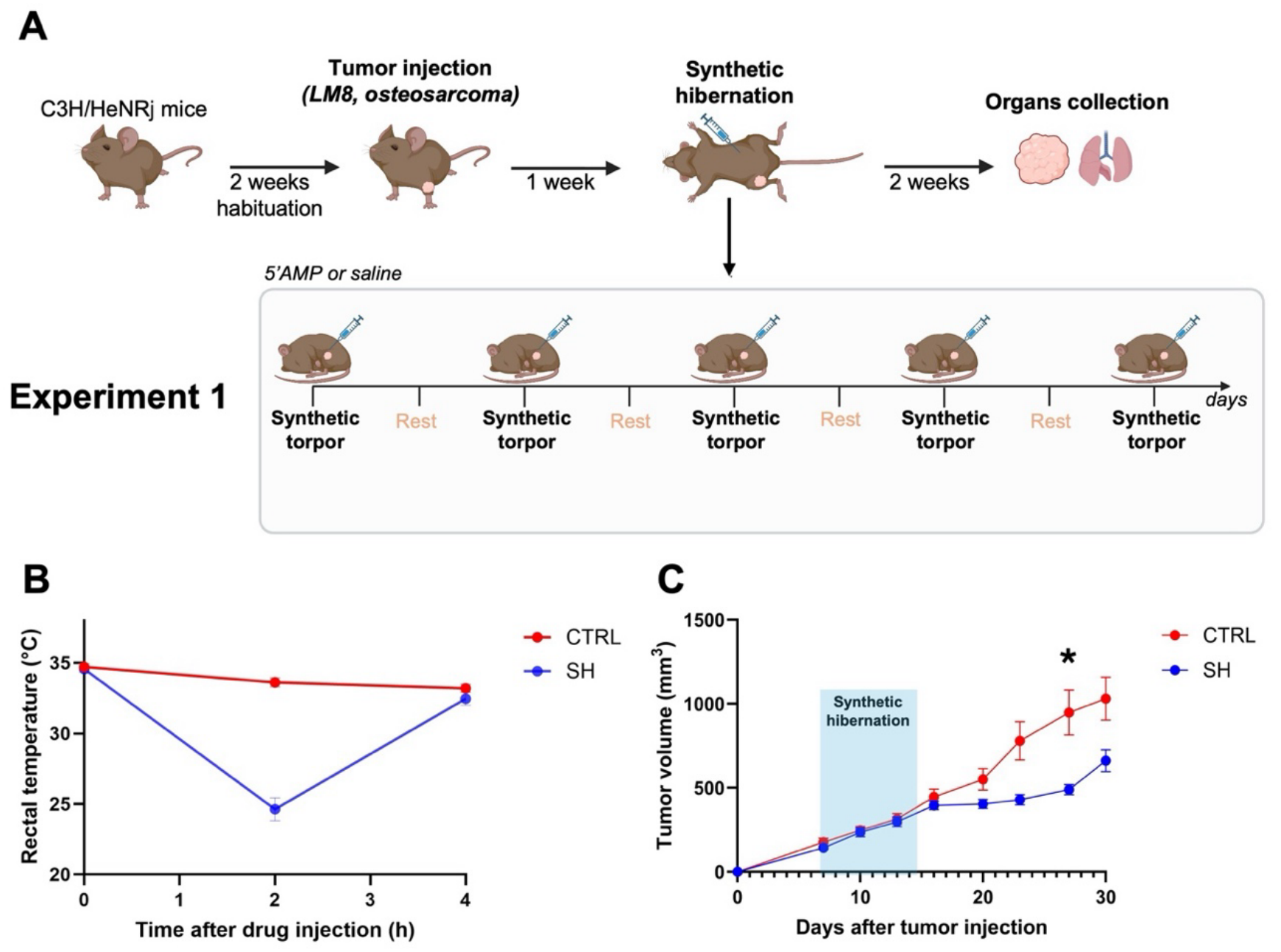
Experimental design, temperature and tumor growth in experiment 1. **A)** Schematic overview of the experimental timeline and repeated synthetic hibernation (SH) protocol. **B)** Rectal temperature profiles after 5′-AMP or saline injection. **C)** Tumor growth curves in SH and CTRL mice. Two-way RM ANOVA: time × treatment, p = 0.0019; treatment, p = 0.0230. Welch’s t-test at day 27, *p = 0.0241. Data are shown as mean ± SEM (n = 9 per group).

### Repeated 5’-AMP induced-SH slows primary tumor growth long after SH

In experiment 1, SH altered tumor growth over time (time × treatment interaction, p = 0.0019; overall treatment effect, p = 0.023) (Fig. 1C). Linear regression of individual trajectories yielded a lower mean growth rate in SH than CTRL mice (SH: 18.82 ± 1.83 mm^3^/day; CTRL: 39.82 ± 5.95 mm^3^/day; Welch’s t-test, p = 0.0076). Apparent doubling time was numerically longer after SH but was not significant in the full dataset (SH: 8.24 ± 1.02 days; CTRL: 6.25 ± 2.00 days). Following exclusion of one CTRL outlier identified by Grubbs’ test, this difference reached statistical significance (Welch’s t-test, p = 0.0044). Tumor volumes were significantly lower in SH mice than in CTRL mice at day 27 (Welch’s t-test, p = 0.0087), during a plateau-like period extending from approximately day 16 to day 27. This difference was no longer significant at day 30, when SH tumors had resumed expansion and fewer CTRL animals remained measurable because they had reached humane endpoints. This late comparison is therefore affected by informative attrition.

### Irradiation retains tumor-growth control during synthetic hibernation

The non-irradiated groups in experiment 2 reproduced the growth-attenuation direction observed in experiment 1. Tumors in SH mice showed a markedly slower expansion than CTRL tumors. Given the differences in tumor volumes between SH and CTRL mice at corresponding time points, growth rates were compared over five consecutive measurements starting from comparable mean tumor volumes (CTRL: days 7–20; SH: days 10–23). Linear regression of individual tumor-volume trajectories showed a significantly lower growth rate in SH mice than in CTRL mice (CTRL: 30.73

± 3.31 mm^3^/day; SH: 12.05 ± 1.31 mm^3^/day; Welch’s t-test, p = 0.0003). Consistently, the apparent time required to double the starting tumor volume was prolonged in SH mice compared with CTRL mice (CTRL: 7.38 ± 0.90 days; SH: 17.42 ± 3.83 days; Welch’s t-test, p = 0.017). At day 27, final tumor volume was significantly reduced in SH, CTRLrad, and SHrad mice compared with CTRL mice (SH vs CTRL, p = 0.018; CTRLrad vs CTRL, p = 0.0089; SHrad vs CTRL, p = 0.0063).

Both irradiated groups (SHrad and CTRLrad) exhibited similar tumor-growth trajectories, characterized by an apparent plateau-like phase. Descriptive fitting of the mean trajectories using a one-phase association model provided a better fit than linear regression in both groups (SHrad: R^2^ = 0.9935 vs. 0.9114, ΔAICc = 11.24; CTRLrad: R^2^ = 0.9811 vs. 0.7828, ΔAICc = 10.10). The estimated rate constants were k = 0.1132 and 0.1884 day^−1^ for SHrad and CTRLrad, respectively. Together, these findings indicate that synthetic hibernation did not measurably compromise irradiation-mediated tumor-growth control (Fig. 2C, 2D).

**Figure 2.**
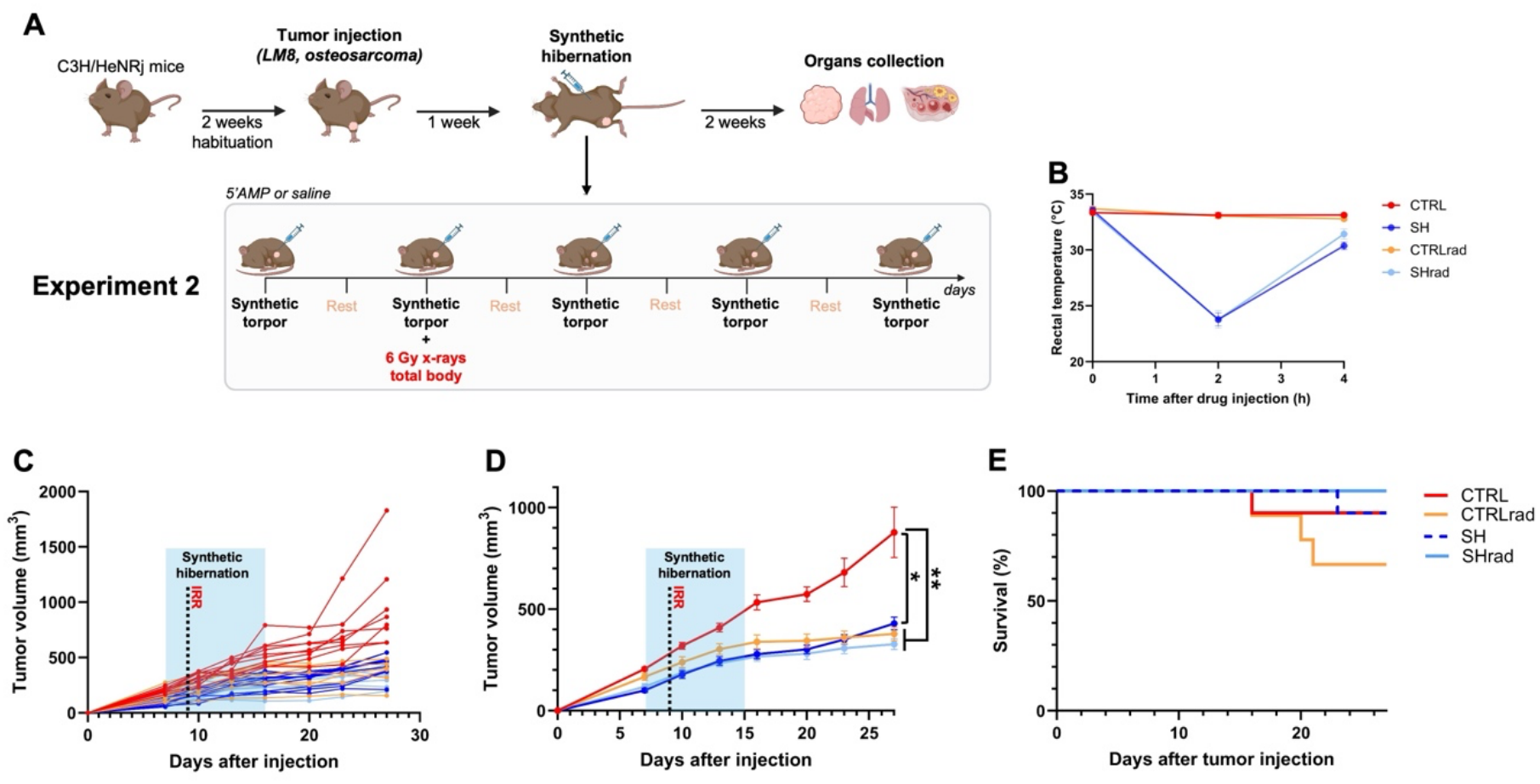
Experimental design, tumor growth, and survival in Experiment 2. **A)** Schematic overview of the experimental timeline, repeated synthetic hibernation (SH) protocol, and irradiation timing. **B)** Rectal temperature profiles after 5′-AMP or saline administration across experimental groups. **C)** Individual tumor growth trajectories. **D)** Mean tumor growth curves. Statistical significance refers to tumor volumes at day 27; comparisons vs CTRL using Welch’s ANOVA followed by Dunnett’s T3 multiple-comparisons test (SH vs CTRL, p = 0.0184; CTRLrad vs CTRL, p = 0.0089; SHrad vs CTRL, p = 0.0063). Data are shown as mean ± SEM. (CTRL and SH, n = 10 per group; CTRLrad and SHrad, n = 9 per group). **E)** Kaplan–Meier survival curves after tumor inoculation.

Kaplan–Meier analysis of time to humane endpoint showed no significant difference among groups. Survival was 100% in SHrad mice (9/9), 90% in CTRL mice (9/10), 90% in SH mice (9/10), and 66.7% in CTRLrad mice (6/9). Although limited by the low number of events, survival was descriptively highest in SHrad mice and lowest in CTRLrad mice (Fig. 2E).

### Synthetic hibernation does not alter lung metastatic burden

In experiment 1, macroscopic counting of superficial nodules in Bouin-fixed lungs showed no difference between SH and CTRL mice (Fig. 3A). In experiment 2, histological quantification of metastatic area across serial lung sections, a method previously used by our group (*32*) likewise showed no difference among CTRL, SH, CTRLrad and SHrad groups (Fig. 3B). These assays therefore provided no evidence that SH altered lung metastatic burden under the conditions and at the endpoints tested.

**Figure 3.**
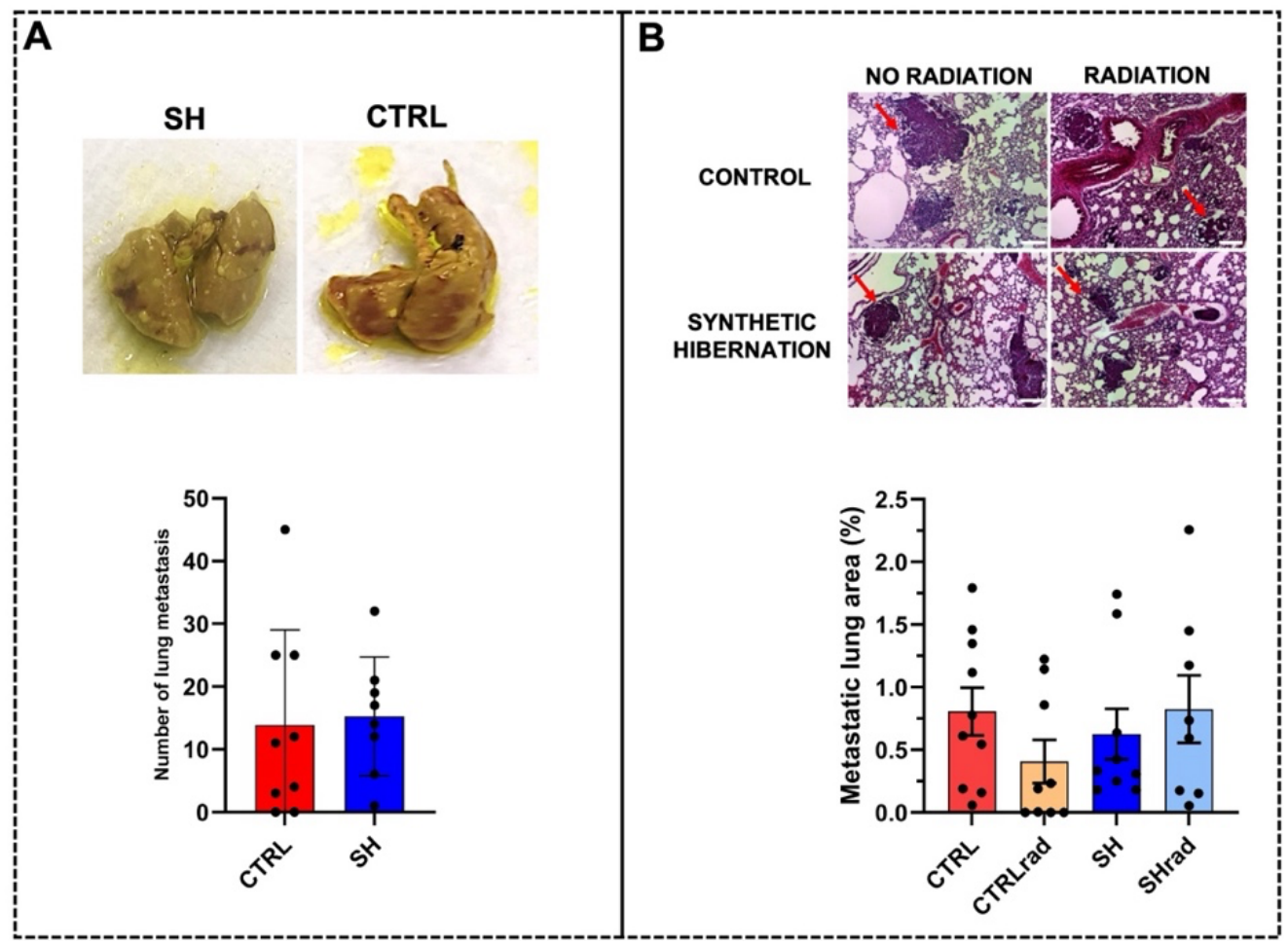
Lung metastatic burden after synthetic hibernation and irradiation. **A)** Experiment 1. Representative images of Bouin-fixed lungs and quantification of superficial metastatic nodules in SH (n = 8) and CTRL (n = 9). **B)** Experiment 2. Representative H&E-stained lung sections across experimental conditions, with metastatic foci indicated by red arrows (scale bars, 200 µm). Quantification shows the percentage of lung area occupied by metastatic tissue. CTRL, n = 10; CTRLrad, n = 9; SH, n = 9; SHrad, n = 8. Data are shown as mean ± SEM; each dot represents one animal.

### Synthetic hibernation attenuates radiation-induced ovarian follicular atresia

Six Gy total body X-ray irradiation increased the proportion of atretic ovarian follicles in normothermic mice (CTRLrad *vs*. CTRL, p = 0.0095). Follicular atresia was lower in SHrad than CTRLrad mice (p = 0.011) and was not detectably different from either non-irradiated group. Importantly, no difference was observed between CTRL and SH mice, indicating that 5′-AMP-induced synthetic hibernation alone did not detectably affect ovarian follicle integrity (Fig. 4). Together these findings indicate that synthetic hibernation mitigates radiation-induced ovarian follicular damage without inducing detectable ovarian toxicity in the absence of irradiation.

**Figure 4.**
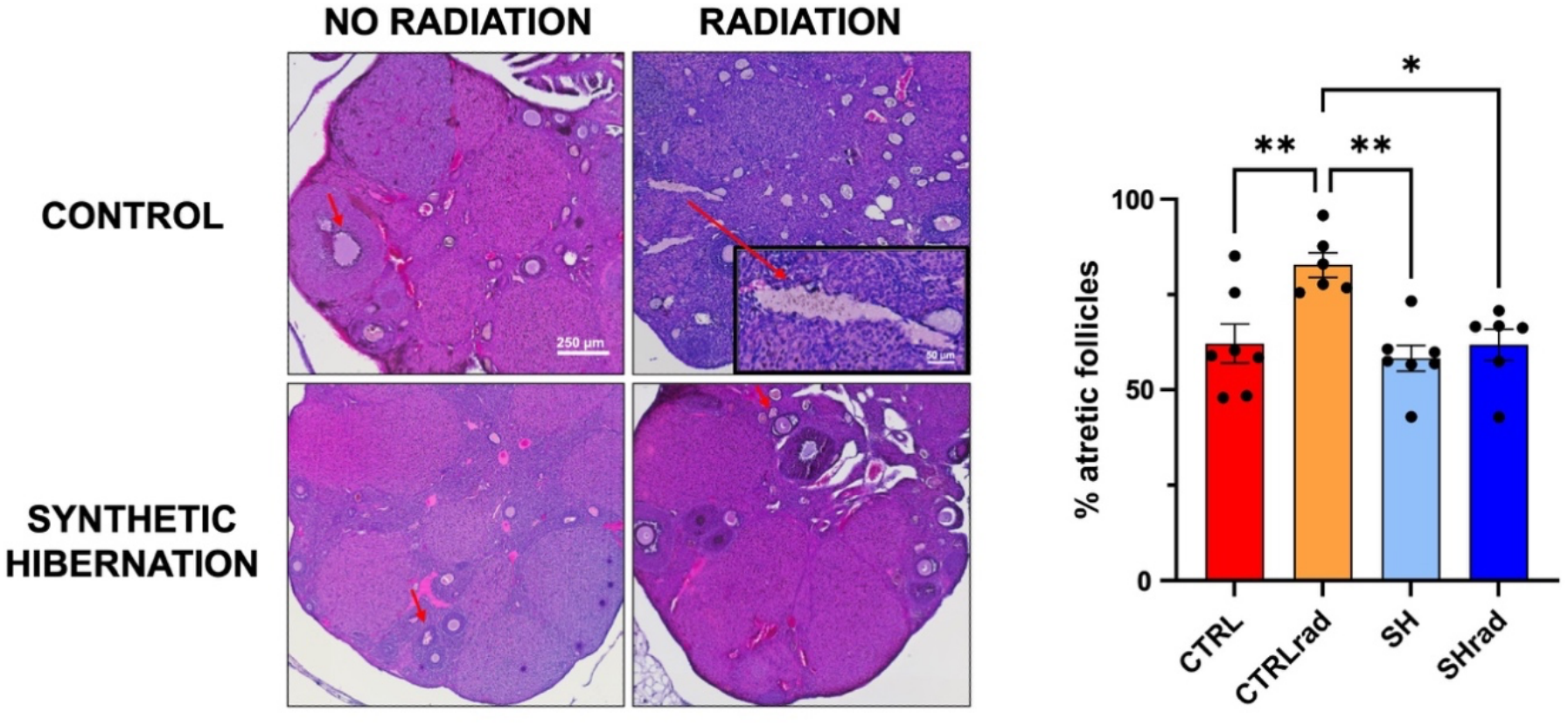
Synthetic hibernation attenuates radiation-induced ovarian follicular atresia. Representative H&E-stained ovarian sections across experimental conditions and quantification of atretic follicles. Arrows indicate atretic follicles (scale bars, 250 and 50 µm). Follicular atresia was expressed as the proportion of atretic follicles over total follicles. Data are shown as mean ± SEM; each dot represents one animal. Statistical comparisons were performed by one-way ANOVA with Tukey’s multiple-comparisons test. *p < 0.05; **p < 0.01.

**Figure 5.**
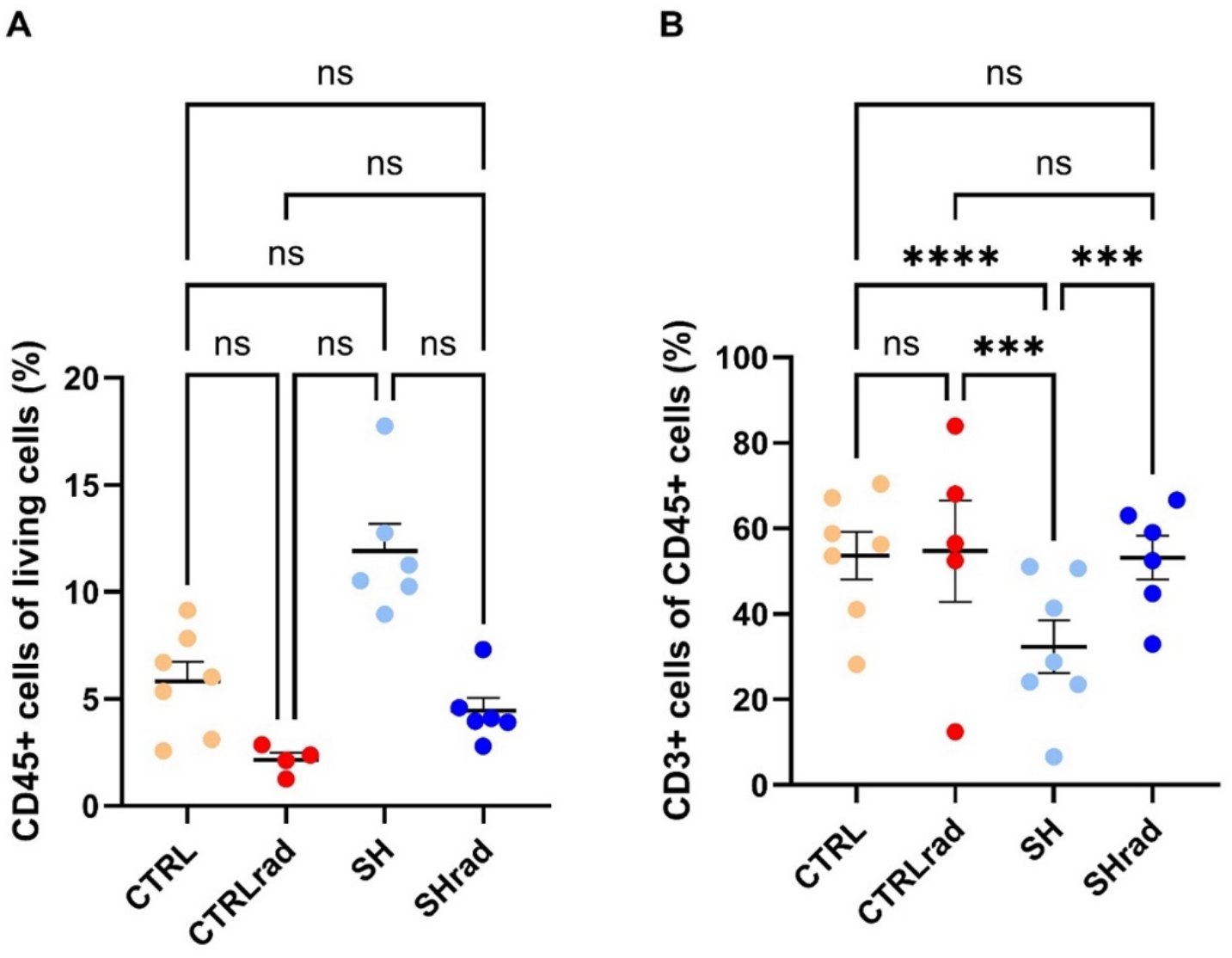
Infiltration of CD45+ leukocytes and CD3+ T-cells in Experiment 2. Flow-cytometry analysis of tumor immune infiltrates. **A)** Percentage of CD45+ leukocytes gated on the total amount of living cells. **B)** Percentage of CD3+ T-cells gated on the total amount of CD45+ cells. Data are shown as mean ± SEM; each dot represents one animal. Statistical comparisons were performed using a two-way ANOVA with Tukey’s multiple-comparisons test (***p < 0.001, ****p < 0.0001).

### Synthetic hibernation is associated with changes in tumor immune infiltration

We next examined the tumor immune microenvironment at late experimental endpoints, i.e. day 27 in Experiment 2, corresponding to 12 days after the last synthetic hibernation bout. Among the lymphoid immune cells analyzed, we observed a tendency for an increased percentage of CD45+ leukocytes (not significant). The percentage of CD3+ T-cells (pan-T-cells) was found significantly decreased in non-irradiated SH tumors compared to all other groups (p < 0.05). Among the pan T-cells, we did not detect significant differences between CD4+ helper or CD8+ cytotoxic T-cells (Fig. S1, supplementary material). Further analyzed immune cell populations also revealed no significant differences and the amount of living cells in the tumor did not change among the groups (Fig. S1 supplementary material). Overall, these data suggest that synthetic hibernation may be associated with increased immune infiltration of leukocytes but a decreased infiltration of pan-T-cells.

### Synthetic hibernation is associated with immune and extracellular matrix transcriptional programmes

To investigate whether SH was associated with persistent tumor transcriptional changes, we performed bulk RNA-seq on endpoint primary tumors from Experiment 2 across the four experimental conditions (n = 3 per group). Weighted gene co-expression network analysis (WGCNA) identified multiple co-expression modules, which were correlated with experimental traits. The MEroyalblue module showed the strongest positive association with SH (r = 0.68, p < 0.001) and minimal association with irradiation, consistent with an SH-associated co-expression program (Fig. 6A). GO analysis of this module revealed enrichment for immune-related biological processes, with 9 of the top 10 enriched GO Biological Process terms related to antigen processing and presentation, particularly MHC class I-mediated peptide presentation (Fig. 6B).

**Figure 6.**
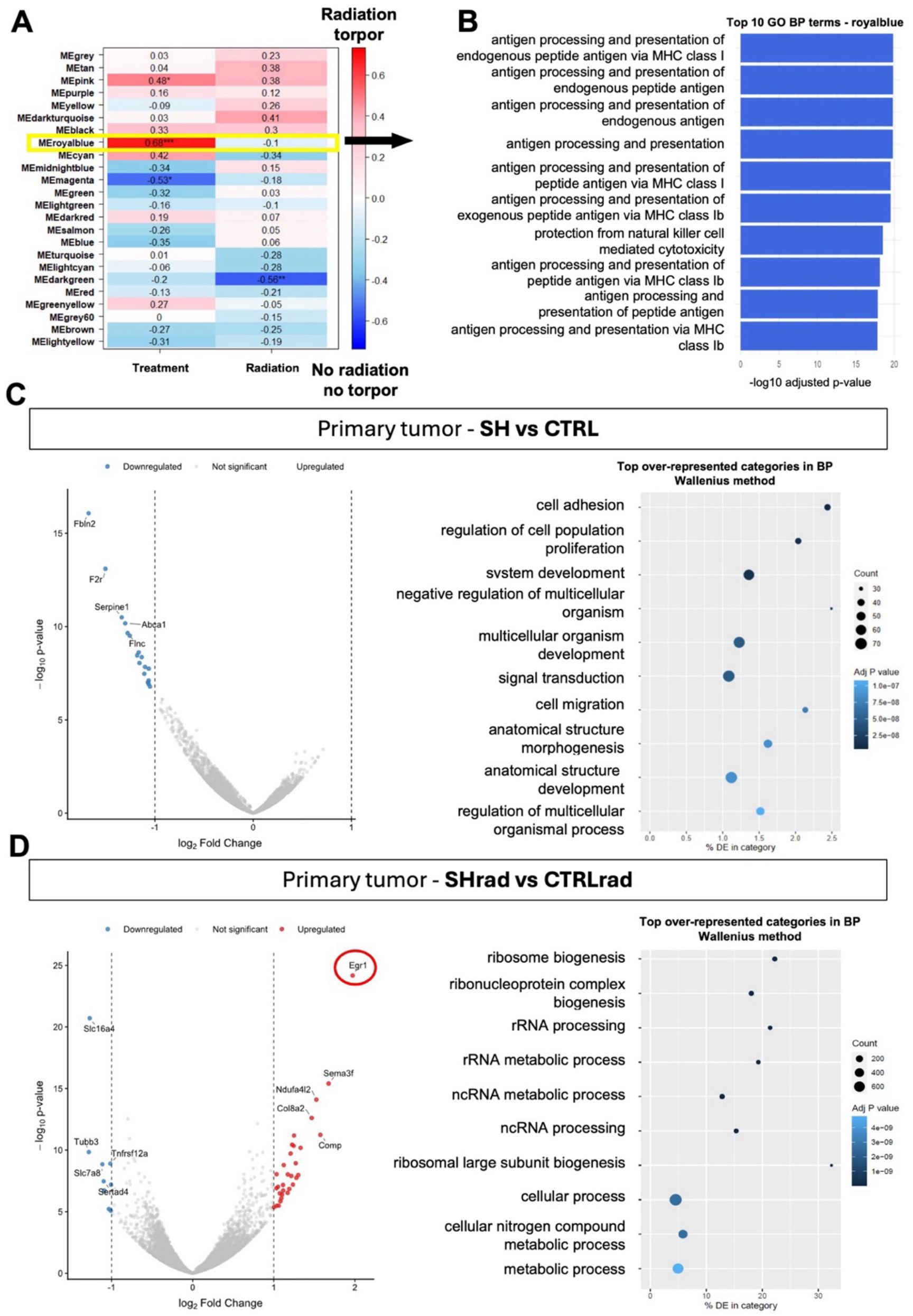
Tumor transcriptomic changes associated with synthetic hibernation. **A)** Heatmap showing how tumor gene co-expression modules identified by WGCNA in Experiment 2 correlate with synthetic hibernation and irradiation. The royalblue module is highlighted and showed the strongest positive correlation with synthetic hibernation (*r* = 0.68). **B)** Top 10 Gene Ontology Biological Process terms enriched in the royalblue module. Bar lengths represent −log_10_ (adjusted *P* value). **C, D)** DESeq2 differential expression analysis of primary tumor cultures, shown by volcano plots and GO Biological Process enrichment: **C)** SH versus CTRL; **D)** SHrad versus CTRLrad. In volcano plots, colored points indicate significant differentially expressed genes; in GO plots, dot size indicates gene count and color indicates adjusted p value.

Pairwise differential expression analysis further showed that SH-associated transcriptional changes differed between non-irradiated and irradiated tumors. In non-irradiated tumors, SH versus CTRL identified a restricted set of differentially expressed genes, all downregulated in SH tumors, including genes linked to adhesion, extracellular-matrix remodeling, and tumor–stroma interactions, such as Fbln2, Fn1, Nid1, Fbn1, Lamb1, Serpine1, and Tgfb2. Corresponding GO enrichment highlighted processes related to cell adhesion, regulation of cell population proliferation, cell migration, signal transduction, and anatomical development/morphogenesis (Fig. 6C). In irradiated tumors, SHrad versus CTRLrad identified a broader response, with 40 differentially expressed genes, including 31 upregulated and 9 downregulated transcripts. Egr1 was the most strongly upregulated gene; GO enrichment highlighted ribosome biogenesis, ribonucleoprotein complex biogenesis, rRNA processing, and ncRNA metabolism (Fig. 6D). Together, these exploratory analyses suggest that SH is associated with persistent tumor transcriptional changes involving immune-related co-expression programs, extracellular-matrix/adhesion genes, and RNA-processing pathways.

## DISCUSSION

This study shows that repeated 5′-AMP-induced synthetic hibernation led to tumor growth suppression that persisted beyond the final torpor bout. When combined with irradiation, SH reduced ovarian follicular injury without a detectable loss of tumor-growth control under the conditions tested.

The attenuation of primary-tumor growth is consistent with both historical and recent observations *(12, 13, 27-31)*. Sarcoma growth in hamsters was reported to pause during hibernation and resume upon arousal (*12, 13*), and Q-neuron-induced hypothermia and hypometabolism have been shown to suppress implanted tumor growth in mice (*30*). The most notable finding in the present experiments was that growth suppression persisted beyond the final torpor bout, although tumors subsequently resumed expansion. This temporal pattern suggests that the effects of SH on tumor growth can outlast the hypothermic phase. Whether this persistence reflects changes within cancer cells, the tumor microenvironment, or systemic signals remains unresolved. Nevertheless, it raises the possibility that intermittent SH could produce periods of tumor-growth suppression without requiring continuous torpor. Establishing whether additional treatment cycles can renew or prolong this effect will be an important next step.

The mechanisms underlying this persistent response remain unclear. Cold-activated brown adipose tissue can compete with tumors for circulating glucose (*33*), illustrating how changes in host metabolism can influence tumor growth. However, this thermogenic response is distinct from the hypometabolic state examined here. Plasma from hibernating common carp has also been reported to suppress tumor growth in mice (*34*), suggesting that circulating factors may contribute to the antitumor effects associated with hibernation. Temperature-dependent reactivation of specific mutant p53 proteins provides another possible link between hypothermia and tumor-growth suppression *(28, 35, 36)*, although its relevance to the present model remains to be established. Persistent changes in cell-cycle regulation, potentially involving epigenetic mechanisms, also warrant investigation.

A second aim of this study was to examine whether SH could reduce radiation injury in healthy tissue while maintaining tumor-growth control. Osteosarcoma-bearing mice therefore received 6 Gy total-body X-ray irradiation during the second torpor bout. The dose was selected to inhibit tumor growth without aiming at complete eradication and was close to the reported LD_50_/_30_ of approximately 7 Gy for this mouse strain. Total-body irradiation allowed tumor response and normal tissue injury be evaluated in the same animals. Among normal tissues, ovaries were selected for analysis because of the high radiosensitivity of ovarian follicles, providing a sensitive endpoint for assessing potential SH-mediated radioprotection (*37, 38*). Tumor-growth trajectories were similar in SHrad and CTRLrad mice, whereas ovarian follicular atresia was significantly lower in SHrad mice and did not differ significantly from non-irradiated controls. This finding extends previous reports of reduced radiation injury during synthetic torpor in rodents (*25, 26*). However, the ovarian endpoint demonstrates reduced histological injury at a single time point, rather than preserved ovarian reserve, fertility.

The absence of a detectable loss of tumor-growth control is encouraging but does not establish unchanged tumor radiosensitivity. Because SH alone also delayed tumor growth, similar trajectories in SHrad and CTRLrad do not exclude partial tumor radioprotection offset by the growth-suppressive effect of SH. Resolving this will require studies using localized irradiation and multiple radiation doses to determine whether SH can improve the therapeutic window and whether normal-tissue protection could permit dose escalation. Tumor oxygenation is another potentially important determinant of radiation response (*39–42*), but was not measured in this study. Whether changes in oxygen supply and demand contribute to the observed response therefore remains unresolved.

Neither lung metastatic burden nor time to humane endpoint differed significantly between groups. Lung assessment included macroscopic nodule counting in the first experiment and histological quantification in the second, with neither approach detecting an effect of SH. The reduction in primary-tumor growth therefore did not translate into a detectable reduction in pulmonary metastases at the endpoints examined. Similarly, the Kaplan–Meier analysis measured time to humane endpoint rather than disease-specific mortality, and the small number of events precludes a conclusion about survival benefit. The absence of a detectable effect on lung metastatic burden may also reflect the timing of metastatic dissemination. Some metastatic lesions may have developed during the post-treatment period, when animals had returned to euthermia, although early dissemination cannot be excluded and the timing of metastatic seeding could not be determined in the present study. The absence of a detectable effect on lung metastases may also provide some insight into the mechanisms underlying tumor-growth suppression. While this finding appears less consistent with a persistent, broadly acting circulating antitumor factor, it does not exclude transient humoral signals acting during the torpor bouts. Indeed, metastatic dissemination and outgrowth may have occurred after the final bout, when such signals were no longer present. However, the timing of metastatic seeding was not determined in this study.

Immune profiling identified a non-significant increase of CD45+ leukocytes among viable cells, however, within these, the amount of CD3+ T-cells (pan-T-cells) was significantly lower in non-irradiated SH tumors. These findings indicate a change in the leukocyte composition, although changes in further investigated populations were only minor. Moreover, samples were collected 12 days after the final torpor bout or 18 days after the irradiation, hence, representing comparably late time points for the investigation of the immune infiltrate after the treatment. The observed differences could therefore have contributed to the persistent growth suppression or reflect its consequences.

Exploratory RNA sequencing of short-term tumor-derived cultures identified additional treatment-associated differences. In non-irradiated samples, SH was associated with lower expression of genes related to extracellular-matrix organization, cell adhesion, and TGF signaling. Co-expression analysis also identified an SH-associated module enriched for antigen processing and presentation, particularly MHC class I-related pathways. In irradiated samples, SH was associated with a broader transcriptional response involving ribosome biogenesis and RNA processing. Notably, *Egr1*, which encodes a transcription factor that can directly regulate Trp53 (43), was the most strongly upregulated gene in SHrad compared with CTRLrad. Given evidence that EGR1 can contribute to radiation-induced growth inhibition and apoptosis in tumor cells and promote apoptosis in osteosarcoma cells under pharmacological treatment (44-46), the higher Egr1 expression in SHrad tumors raises the possibility that EGR1 contributes to growth restraint in this group. These findings identify candidate pathways for further investigation, but do not establish increased tumor immunogenicity or functional inhibition of the processes identified by Gene Ontology analysis. Because RNA was extracted after short-term culture, these profiles should not be interpreted as a direct representation of the intact tumor microenvironment. Their contribution to the persistent growth response requires functional validation.

Several limitations should guide further work. The effects of 5′-AMP, reduced body temperature, and metabolic suppression were not experimentally separated, preventing attribution of the antitumor response to a single component of the protocol. Only one syngeneic tumor model, one total-body radiation dose, and female mice were studied. Although the female cohort enabled assessment of ovarian injury, sex-dependent effects on tumor growth and treatment response remain unresolved. Additional tumor models, independent methods of torpor induction, and longer follow-up will be needed to establish the reproducibility, duration, and broader relevance of these findings. In summary, synthetic hibernation suppressed osteosarcoma growth beyond the final torpor bout and reduced ovarian radiation injury, with no detectable loss of tumor-growth control during combined treatment. These findings support further investigation of SH as an antitumor intervention and as a potential adjunct to radiotherapy.

## MATERIALS AND METHODS

### Ethical Approval

All animal procedures were performed in accordance with the German Federal Animal Welfare Act and were approved by the Hessen Animal Ethics Committee (Project License DA 17/2005, Regierungspräsidium Darmstadt, Germany).

### Animals and Experimental Design

A total of 56 female C3H/HeNRj mice (Janvier Labs, Saint-Berthevin Cedex, France), 11 weeks old upon arrival, were included in two independent experiments. Animals were housed in individually ventilated cages with bedding and environmental enrichment under standard laboratory conditions, with ad libitum access to standard food (ssniff Spezialdiäten GmbH, Soest, Germany) and water. The animal room was maintained at an ambient temperature of 22.0 ± 1.0 °C and 55% relative humidity, under a 12 h:12 h light–dark cycle, with lights on from 08:00 to 20:00. Cages, bedding, and nesting material were changed once weekly. After one week of acclimatization, mice were provided with a high-calorie diet to limit weight loss potentially associated with the synthetic hibernation and/or irradiation procedures. One week later, mice were randomly assigned to the experimental groups.

*Experiment 1* was designed to evaluate the effect of synthetic hibernation on tumor growth. Mice were allocated to synthetic hibernation (SH, n = 9) or normothermic control conditions (CTRL, n = 9) (Figure 1A). *Experiment 2* was designed to assess whether the radioprotective state induced by synthetic hibernation could interfere with irradiation-mediated tumor control. Mice were randomly assigned to four experimental groups: synthetic hibernation with irradiation (SHrad, n = 9), synthetic hibernation without irradiation (SH, n = 10), irradiated controls (CTRLrad, n = 9), and non-irradiated controls (CTRL, n = 10) (Fig. 2A).

### Osteosarcoma Model and Tumor Monitoring

Dunn osteosarcoma LM8 cells, derived from C3H/He mice and obtained from the RIKEN BioResource Center, were used to establish a syngeneic osteosarcoma model. Mice were subcutaneously injected in the right hind limb with 1 × 10^6 LM8 cells suspended in 20 µL phosphate-buffered saline. Tumor growth was monitored twice weekly, starting one week after tumor implantation and continuing until sacrifice. Tumor length and width were measured with a caliper and tumor depth was estimated using a standardized measurement that included the thickness of the tumor-bearing hind limb. All measurements were performed by the same investigator using the same procedure throughout the experiment. Since tumors were approximately ellipsoid in shape, tumor volume was calculated as:

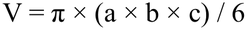

where *a, b* and *c* representing tumor length, width, and depth, respectively.

Animals were euthanized when humane endpoints were reached, including tumor diameter exceeding 1.5 cm, marked body weight loss, poor general condition, abnormal coat appearance, or hunched posture. For animals euthanized before the experimental endpoint, subsequent tumor-volume values were treated as missing and were not carried forward.

To characterize tumor-growth kinetics, linear regression of individual tumor-volume trajectories was used to estimate growth rates (mm^3^/day). Apparent doubling time was calculated as the ratio of the initial tumor volume within the analyzed interval to the corresponding linear growth rate. In Experiment 2, SH and CTRL growth rates were compared over five consecutive measurements starting from comparable mean tumor volumes (CTRL: days 7–20; SH: days 10–23).

To further characterize growth patterns in irradiated tumors, mean tumor-volume trajectories from SHrad and CTRLrad mice were descriptively fitted using a one-phase association model:

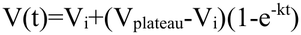

where V_i_ represents the fitted initial tumor volume, V_plateau_ the estimated asymptotic volume, and k the rate constant (day^−1^). Time was expressed relative to the first measurement included in the fit (day 7 after inoculation). Model fit was compared with linear regression using the corrected Akaike information criterion (AICc), with ΔAICc defined as AICc_(linear)_ − AICc_(plateau)_. Positive ΔAICc values indicate a better fit of the plateau model.

### Synthetic Hibernation Induction

One week after tumor implantation, mice assigned to the synthetic hibernation groups received an intraperitoneal injection of 5′-AMP (900 mg/kg; Sigma-Aldrich, St. Louis, MO, USA) to induce a hibernation-like state. Control mice received an equivalent injection of saline. Injections were performed at room temperature (~21°C ± 2°C) during the inactive light phase of the circadian cycle. After injection, mice were placed individually in small round chambers. Cutaneous body temperature was monitored using an infrared thermography camera (FLIR-E6390, FLIR Systems), and rectal temperature was measured at 0, 2h and 4h post injection using a rectal probe. Once mice had returned to normothermia (approximately 4 hours after drug injection), they were placed back in their home cages. Control mice were handled according to the same timing as the synthetic hibernation groups. This procedure was repeated every other day for a total of five synthetic torpor bouts (Figure 1A, 2A).

### Irradiation Protocol

In *Experiment 2*, irradiation was performed during the second day of synthetic hibernation session (Figure 2A). Two hours after 5′-AMP or saline injection, corresponding to deep torpor in the synthetic hibernation groups, mice were anesthetized with isoflurane (3% for induction and 1.5% for maintenance) and placed in a Plexiglas box supplied with an isoflurane–air mixture. Ophthalmic ointment was applied to prevent corneal drying and reduce radiation-related ocular damage. Mice assigned to the irradiated groups received 6 Gy total-body X-ray irradiation at a dose rate of 2 Gy/min using an in-house X-ray generator (Isovolt DS1, Seifert, Ahrensberg, Germany) operated at 250 kVp and 16 mA, with 7 mm beryllium, 1 mm aluminum, and 1 mm copper filtering. Non-irradiated animals underwent the same procedure without radiation exposure. After irradiation or sham irradiation, isoflurane was replaced with fresh air. The hypothermic animals were allowed to continue their torpor bouts, whereas control animals were placed on a heating pad until they recovered a normothermic conscious state.

### Euthanasia and Tissue Collection

Thirty days after tumor inoculation in Experiment 1 and 27 days after tumor inoculation in Experiment 2, mice were deeply anesthetized with isoflurane and euthanized by cervical dislocation. Following tumor collection, the thoracic cavity was opened, and lungs and ovaries were then collected for histological, immunological and/or RNA-seq analyses.

### Histology and Image Analysis

Lung metastatic burden was assessed at different levels of resolution in the two experiments. In Experiment 1, lungs were immersed in Bouin’s solution to visualize superficial metastatic nodules. In Experiment 2, metastases were evaluated by histological sectioning and quantitative image analysis, enabling assessment within the lung parenchyma. Ovarian analysis was performed only in Experiment 2, which included irradiation, using the ovary as a radiosensitive organ to assess normal-tissue radioprotection.

For histological analyses in Experiment 2, lungs and ovaries were fixed in 4% paraformaldehyde for 24 h and then stored in 70% ethanol at 4 °C until paraffin embedding. To obtain representative sampling across each organ, lungs and ovaries were sectioned sequentially at 500 µm and 200 µm intervals, respectively. At each interval, 5 µm-thick sections were obtained using a microtome (Leica RM2125, Leica Microsystems, Germany). Sections were stained with hematoxylin and eosin (H&E) to assess tissue morphology. Lung metastases and ovarian follicular atresia were quantified using QuPath software. In lung sections, metastatic areas were manually annotated and expressed as a percentage of total lung area: metastatic area (%) = (metastatic area / total lung area) × 100. Metastatic burden was assessed at increasing tissue depths (500–2000 µm), and values were averaged for each animal to obtain the mean percentage of lung metastatic area. In ovarian sections, the total number of follicles and the number of atretic follicles were counted at 600 and 800 µm depth, and follicular atresia was expressed as the proportion of atretic follicles over total follicles.

### Flow cytometry analysis of immune cells infiltrating the tumor

To obtain single cell suspensions, the isolated tumors were manually disrupted using tissue strainers (40 or 100 µm; Greiner Bio-One). A subsequent lysis of red blood cells was performed for 5 minutes in a buffer containing ammonium chloride, potassium hydrogencarbonate and EDTA disodium salt. Tumors were treated with a tumor dissociation kit (Miltenyi Biotec) and subjected to a dead cell removal (Miltenyi Biotec). Subsequently, cell suspension was washed in phosphate-buffered saline, centrifuged, and resuspended in a medium consisting of Roswell Park Memorial Institute 1640 Medium (RPMI 1640), fetal calf serum, Minimum Essential Non-Essential Amino Acides (MENEAA), Pen/Strep, L-Glutamin, and sodium pyruvate. The number of viable cells per mL was counted by using 0.4% trypan blue solution (ThermoFisher Scientific) and an automated cell counter (TC20, Bio-Rad Laboratories). Subsequently, 2×10^6^ cells per sample were separated for each staining panel. After blocking the cells in a buffer containing mouse serum, fetal calf serum and optional Fc-Block (BD Bioscience), cell surface proteins of interest were labeled with respective antibodies according to manufacturer’s instructions. The immune cell distribution was measured with a CytoFLEX S flow cytometer (Beckman Coulter) and data were analyzed with the software Cytexpert v2.4 (Beckman Coulter). Debris were further excluded applying FSC/SSC channels. Doublettes were excluded plotting FSC A vs. FSC H. Living cells were defined by Zombie Aqua exclusion. The population of living cells was used to gate for CD45+ leukocytes. Further cell populations were defined as follows: CD3+ pan T-cells (Aqua-, CD45+, CD3+) CD4+ helper T-cells (Aqua-, CD45+, CD3+, CD8-, CD4+), CD8+ cytotoxic T-cells (Aqua-, CD45+, CD3+, CD8+, CD4-), NK cells (Aqua-, CD45+, CD3-, CD49b+), B19+ B-cells (Aqua-, CD45+, CD19+). Antibodies are listed in the Supplementary material (Table S1).

### RNA sequencing

Following animal dissection, tumors were enzymatically dissociated for 30 min at 37 °C in 5 mL of DMEM containing 0.1% (w/v) collagenase I and 3 mM of CaCl2. The resulting cell suspensions were passed through a 100-µm cell strainer and cultured in T75 flasks for 2–3 days. Subsequently, the cells were harvested, and total RNA was isolated using the NucleoSpin RNA Plus kit (Macherey-Nagel) according to the manufacturer’s instructions. RNA sequencing was performed by GENEWIZ (Germany) on an Illumina NovaSeq 6000 platform, generating approximately 20 million paired-end 150-bp reads per sample.

### Bioinformatic analyses

Bioinformatic analysis of the RNA-sequencing data was partly performed on the European Galaxy server (*47*). Read quality was assessed and adapter sequences were removed using fastp. The processed reads were subsequently aligned to the Mus musculus reference genome GRCm39 (mm39) using RNA STAR. Gene-level read counts were generated using featureCounts and used as input for differential gene expression analysis with DESeq2. Gene Ontology enrichment analysis was performed using goseq, and volcano plots were generated using the Galaxy Volcano Plot tool.

Weighted gene co-expression network analysis (WGCNA) was performed in R using the WGCNA and DESeq2 packages. The network construction was performed with cell cultures derived from primary tumors and circulating tumor cells. Genes and samples were assessed using the goodSamplesGenes function, hierarchical clustering and principal component analysis. Genes with at least 15 reads in a minimum of 75% of the samples were retained, resulting in 13,284 genes, and count data were variance-stabilized using the DESeq2 vst function. Candidate soft-thresholding powers were evaluated using pickSoftThreshold, and a power of 12 was selected for network construction. Co-expression modules were identified using blockwiseModules with a maximum block size of 15,000 genes, a signed topological overlap matrix, a module-merging threshold of 0.25, and a random seed of 1,234. Module eigengenes were correlated with binarized treatment (SH) and irradiation using Pearson correlation, and corresponding P values were calculated using the corPvalueStudent function. Modules associated with at least one experimental trait at an unadjusted P value < 0.05 were subjected to Gene Ontology enrichment analysis using clusterProfiler and org.Mm.eg.db. Enrichment analyses were performed separately for Biological Process, Molecular Function, and Cellular Component terms using Ensembl gene identifiers, and P values were adjusted using the Benjamini–Hochberg method; terms with an adjusted P value < 0.05 were considered significantly enriched.

### Statistical Analysis

Data are presented as mean ± SEM. Statistical analyses were performed using GraphPad Prism 6.00. Individual animals were considered the experimental units. Two-group comparisons were performed using unpaired two-tailed Welch’s t-tests. Multiple-group comparisons at single time points or endpoints were performed using one-way ANOVA followed by Tukey’s multiple-comparisons test when variances were homogeneous. When variance heterogeneity was detected, Brown-Forsythe/Welch ANOVA followed by Dunnett’s T3 multiple-comparisons test was used. Longitudinal tumor growth in experiment 1 was analyzed using two-way repeated-measures ANOVA, with time and treatment as factors; Geisser–Greenhouse correction was applied. Tumor growth rates were calculated by linear regression of individual tumor-volume trajectories using measured tumor volumes. Survival was analyzed using Kaplan–Meier curves and log-rank tests. Statistical significance was set at p ≤ 0.05.

For flow cytometry analysis, statistical comparisons were performed using a two-way ANOVA with Tukey’s multiple-comparisons test. Outliers were identified using Tukey’s fences with a k of 1.5.

## Acknowledgments

The authors acknowledge the use of artificial intelligence (AI)-based tools for language refinement and for improving the clarity and readability of the manuscript.

The authors also gratefully acknowledge the contributions of Nicola Loi and Elisabeth van der Bijl, whose work and support helped facilitate the completion of this manuscript.

