## Supplementary for "Synthetic Hibernation Suppresses Osteosarcoma Growth and Protects Normal Tissue from Radiation"

**This PDF file includes:**

Figs. S1
Tables S1

### Supplementary Materials

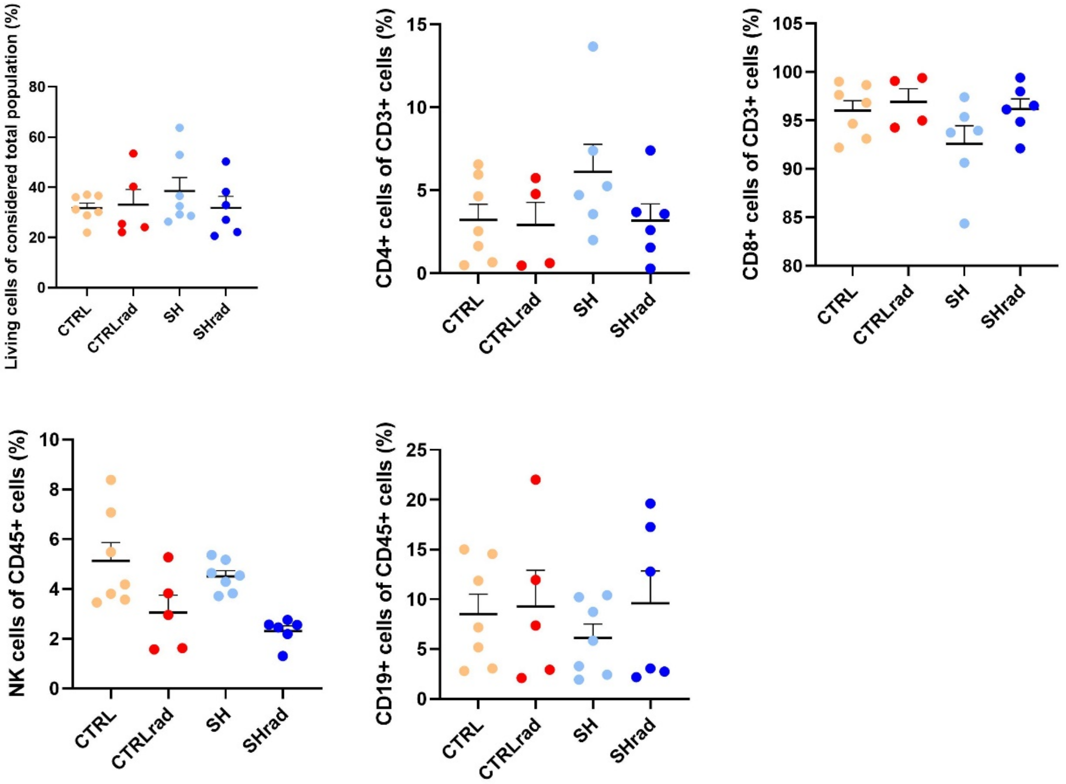


**Fig. S1. Infiltration of CD4+ helper and CD8+ cytotoxic T-cells, NK cells, B19+ B-cells as well as the all living cells in Experiment 2.** Flow-cytometry analysis of tumor immune infiltrates. Data are shown as mean ± SEM; each dot represents one animal. Statistical comparisons were performed using a two-way ANOVA with Tukey's multiple-comparisons test. No significant differences were detected.

| Fluorophor | Marker | Species | Clone | Company | Isotype |
|---|---|---|---|---|---|
| V450 | CD3 | Syr.Ham./anti-mouse | 500A2 | BD Biosciences | IgG2 K |
| Zombie Aqua | Live/Dead | N/A | N/A | BioLegend | N/A |
| Brilliant Violet 605 | CD8a | Rat/anti-mouse | 53-6.7 | BioLegend | IgG2a K |
| APC | CD49b | Rat/anti-mouse | DX5 | BioLegend | IgM K |
| APC-Cy7 | CD19 | Rat/anti-mouse | 1D3 | BD Biosciences | IgG2a K |
| FITC | CD4 | Rat/anti-mouse | GK1.5 | BD Biosciences | IgG2b K |
| PerCP-Cy5.5 | CD45.2 | Mouse/anti-mouse | 104 | ThermoFisher | IgG2b K |

**Table S1. Antibodies for the flow cytometry analysis of infiltrating immune cells.**
